# The neural representation of missing speech and the influence of prior knowledge on cortical fidelity and latency

**DOI:** 10.1101/251793

**Authors:** Francisco Cervantes Constantino, Jonathan Z. Simon

## Abstract

In naturally noisy listening conditions, for example at a cocktail party, noise disruptions may completely mask significant parts of a sentence, and yet listeners may still perceive the missing speech as being present. Here we demonstrate that dynamic speech-related auditory cortical activity, as measured by magnetoencephalography (MEG), which can ordinarily be used to directly reconstruct to the physical speech stimulus, can also be used to “reconstruct” acoustically missing speech. The extent to which this occurs depends on the extent that listeners are familiar with the missing speech, which is consistent with this neural activity being a dynamic representation of perceived speech even if acoustically absence. Our findings are two-fold: first, we find that when the speech is entirely acoustically absent, the acoustically absent speech can still be reconstructed with performance up to 25% of that of acoustically present speech without noise; and second, that this same expertise facilitates faster processing of natural speech by approximately 5 ms. Both effects disappear when listeners have no or very little prior experience with a given sentence. Our results suggest adaptive mechanisms of consolidation of detailed representations about speech, and the enabling of strong expectations this entails, as identifiable factors assisting automatic speech restoration over ecologically relevant timescales.

## 1 Introduction

The ability to correctly interpret speech despite disruptions masking a conversation is a hallmark of communication (Cherry, 1953). In many cases, contextual knowledge poses clear informational advantages for a listener, so as to successfully disengage the masker and restore the intended template signal (Shahin et al., 2009; Riecke et al., 2012; van Wassenhove and Schroeder, 2012; Leonard et al., 2016; Cervantes Constantino and Simon, 2017). Relevant information is available from multimodal sources and/or low-level auditory and higher-level linguistic analyses, although it remains unclear how and which factors are most effective in assisting speech restoration under natural conditions. For instance, while cortical network activity profiles have been identified that are consistent with phonemic restoration (the effect where absent phonemes in a signal may nonetheless be heard (Samuel, 1996, 1981)) in binary semantic decision tasks (Leonard et al., 2016), the factors that bias into one or the other of two perceptual alternatives remain unclear. There is evidence that such restorative processes may be influenced by contributions from audiovisual integration cues (Crosse et al., 2016), lexical priming (Sohoglu et al., 2012), and within the auditory domain, by predictive template matching (SanMiguel et al., 2013) or even intentional expectations about temporal patterns in sound (Nozaradan et al., 2011; Tal et al., 2017).

In order to affect ongoing speech percepts, the potential outcomes from these mechanisms must be readily accessible before and during missing auditory input. These type of contributions might entail (i) generation of a provisional template of the forthcoming speech, (ii) that the template be stored in a compatible format with the internal representation of ongoing sound, and (iii) that they are later subject to point-wise matching – in what has been termed the *zip metaphor* (Bendixen et al., 2014; Grimm and Schröger, 2007; Tavano et al., 2012). In addition, the contribution by such putative mechanisms in enhancing the neural representation of speech may allow a speed up of cortical processing during integration (van Wassenhove et al., 2005).

Here we test how a string of natural speech tokens, spanning several words, may be represented cortically, even if entirely removed and replaced by stationary masking noise—under different levels of informational gain provided by prior knowledge of the masked elements. We use the fact that the low-frequency envelope of speech (i.e., spanning several words) indexes the acoustic signal’s slow changes over time and is known to phase-lock neural activity in auditory cortex, as measured by magnetoencephalography (MEG) and electroencephalography (EEG) (Di Liberto et al., 2015; Ding and Simon, 2012a; Giraud et al., 2000; Zion Golumbic et al., 2013). Because of its timescale, the low-frequency envelope of speech typically reveals attributes such as the patterns of syllabic lengths and loudness changes, as well as prosodic information including intonation, rhythm and stress cues. We hypothesize that by repeating the strings of speech tokens, and controlling for the extent of repetition, it becomes possible to manipulate listeners’ ability to develop detailed predictions about forthcoming elements in these long sentences. More repetitions would allow the generation of a better template for those tokens, to serve for a point-wise matching when later, spontaneous maskers disrupt the same string of tokens. Availability of a temporally-detailed template of the absent speech may allow the missing speech to be decoded from cortical signals representing a token, despite the acoustic absence of the speech itself. Furthermore, because the template would be formed in advance, we also addressed the possibility that cortical representations of highly repeated speech stimuli may be facilitated in terms of processing time for those same speech tokens, even when *not* absent.

To address these hypotheses, we employ complementary systems-based neural analysis methods. In one case, we analyze neural responses in a way that allows reconstruction of a stimulus speech envelope (Mesgarani, 2014), an approach that has been successfully applied in auditory electrophysiology (Mesgarani et al., 2009; Ramirez et al., 2011), EEG/MEG (Ding and Simon, 2012b; O’Sullivan et al., 2015), electrocorticography (Leonard et al., 2016; Pasley et al., 2012), and fMRI (Naselaris et al., 2011). The performance of this decoding method allows a quantitative assessment of the extent to which prior knowledge of absent speech may enhance endogenous representations involved in its perceptual restoration. In the other case we instead use the stimulus speech envelope to estimate the neural response (Di Liberto et al., 2015; Ding and Simon, 2012a), under normal (non-absent) speech conditions. In this forward model case, we analyze cortical latencies involved in natural speech processing under different prior knowledge conditions. The possibility of reduced cortical latencies is of particular interest since faster processing has been observed in situations where additional context facilitates integration of incoming speech (van Wassenhove et al., 2005; van Wassenhove and Schroeder, 2012). Additionally, similar task-related cortical plasticity changes in stimulus-response mappings are often observed at the neuronal level (David et al., 2012; Fritz et al., 2003) and represent a potential biophysical basis for restorative mechanisms given the present task demands.

We provide evidence that the speech temporal envelope is better reconstructed when listeners have obtained more knowledge about a particular speech sequence, and, critically, that this effect applies even in the case where the speech itself is absent, having been replaced entirely with noise. The data also show that cortical latencies in the processing of clean speech can be reduced by several milliseconds when the listener has obtained more knowledge about that particular speech sequence. Overall, the results suggest that the formation of online templates representing low-level features of frequently experienced speech may facilitate more efficient neural representations, both by means of faster encoding and by improved access to endogenous dynamic neural speech representations, time-locked to expected but missing speech, thus assisting its restoration.

## 2 Materials and methods

### 2.1 Participants

35 experimental subjects (19 women, 21.3 ± 2.9 years of age [mean ± SD]), with no history of neurological disorder or metal implants, participated in the study. Data from one additional subject was not included, due to excessive artifacts caused by a poor fit with the MEG helmet. Each subject received monetary compensation proportional to the study duration (approximately 1.5 hours). This study was carried out in accordance with the recommendations of the UMCP Institutional Review Board with written informed consent from all subjects. All subjects gave written informed consent in accordance with the Declaration of Helsinki. The protocol was approved by the UMCP Institutional Review Board.

### 2.2 Stimuli and experimental design

Sound stimuli were prepared with the MATLAB^®^ software package (MathWorks, Natick, United States) at a sampling rate of 22.05 KHz, and consisted of a recorded poem (“A Visit from St. Nicholas”, Moore or Livingston, 1823) obtained from an online archive <http://archive.org/details/AVisitFromSt.Nicholas-ByClementClarkeMoore-NarratedByGrantRaymond>. Each of the fourteen verses (each verse being a quatrain of four lines) in the poem were separated and used as individual stimuli. Silence intervals (gaps) in the narration were reduced to approximately equalize stimuli durations (range: 13.1 – 13.6 s). Four stimulus blocks were presented in total, each containing 64 stimuli (i.e., 256 lines), with some stimuli repeated multiple times. For the first block, a verse from the first half of the poem was chosen as a ‘High’ frequency stimulus, repeated for half of the cases (32/64); similarly, different verses were chosen as ‘Medium’ and ‘Low’ frequency stimuli, which were repeated for a quarter (16/64) and an eighth (8/64) of the cases, respectively. The remainder of the block was filled with ‘Control’ stimuli, namely the four remaining verses presented either 1, 2 or 4 times within the block. Stimuli were randomized in order and concatenated in time. For the second block the same procedure was followed using material from the second half of the poem. Blocks 3 and 4 consisted of the same stimuli used as in 1 and 2 respectively, but with a different randomized order and different placement of noise probes (see below). The procedure was recreated with different randomizations for each subject, resulting in a total of 35 different stimulus sets of about 1 hour each in total duration. Importantly, though, the usage of particular stimuli at a given repetition level was controlled across participants, resulting in seven groups of 5 listeners each that underwent the same ‘High’, ‘Medium’, ‘Low’, and ‘Control’ stimuli selection.

For each stimuli, 2–4 spectrally-matched noise probes of 800 ms duration each were applied at pseudo-random times with a minimum 2.5 s between probe onsets. Noise onset times were selected from a pool of values indicating articulation onset times (e.g. syllables), obtained as the envelope rising slope maxima. Thus 768 noise probe samples were presented per experiment, and each was individually constructed by randomizing phase values across the specific frequency-domain phase information contained in the underlying speech stimulus that would have occurred at the same time as the masker noise, yielding a noise with equal spectral amplitude characteristics (Prichard and Theiler, 1994). The original speech content occurring during the same time was removed entirely and substituted with this spectrally-matched noise, at a power signal level matching that of the excised clean original. Subjects listened to the speech sounds while watching a silent film. To ensure attention to the auditory stimulus, after each probe, they were instructed to report via a button press whether they understood what the speaker meant to say during the noise. The button presses are not analyzed here.

### 2.3 Data recording

We recorded neural responses using MEG, a non-invasive neuroimaging technique well-suited to measure dynamical neural activity from human cortex, and especially from auditory cortical areas. Such recordings typically demonstrate time-locked neural responses to speech low frequency modulations, especially of the acoustic energy envelope, with remarkable temporal fidelity (Ding and Simon, 2012a). MEG data were collected with a 160-channel system (Kanazawa Technology Institute, Kanazawa, Japan) inside a magnetically-shielded room (Vacuumschmelze GmbH & Co. KG, Hanau, Germany). Sensors (15.5 mm diameter) were uniformly distributed inside a liquid-He Dewar, spaced ∼25 mm apart. Sensors were configured as first-order axial gradiometers with 50 mm separation and sensitivity > 5 fT·Hz^-1/2^ in the white noise region (> 1 KHz). Three of the 160 sensors were magnetometers employed as environment reference channels. A 1 Hz high-pass filter, 200 Hz low-pass filter, and 60 Hz notch filter were applied before sampling at 1 KHz. Participants lay supine inside the magnetically shielded room under soft lighting, and were asked to minimize movement, particularly of the head.

### 2.4 Data processing

#### Pre-processing and sensor rejection

The time series of raw recordings from the MEG sensor array were be submitted to a fast implementation of independent component analysis (Hyvärinen, 1999), from which two independent components were selected for their maximal proportion of broadband (0-500 Hz) power (because of the ∼1/f power spectrum of typical neural MEG signals, these components are dominated by non-neural artifacts). These independent components, combined with the physical reference channels, were treated as environmental noise sources arising from unwanted electrical signals not related to brain activity of interest, and were removed using time-shifted principal component analysis (TS-PCA)(de Cheveigné and Simon, 2007). Sensor-specific sources of signals unrelated to brain activity were reduced by sensor noise suppression (SNS)(de Cheveigné and Simon, 2008a).

### 2.5 Data analysis

To analyze low-frequency cortical activity, recordings were bandpass filtered between 1 and 8 Hz with an order-2 Butterworth filter, with correction for the group delay. A blind source separation technique, Denoising Source Separation (DSS)(de Cheveigné and Simon, 2008b), was used to construct components (virtual channels constructed of linear combinations of the sensor channels), ranked in order of their trial-to-trial reproducibility, and used as described below.

#### 2.5.1 Stimulus reconstruction

The ability to reconstruct the speech stimulus envelope from recorded neural responses was used to measure the dynamical cortical representation of perceived speech. The first three DSS components (i.e. with highest reproducibility) were used to train an optimal linear decoder, designed to reconstruct the envelope of the stimulus responsible for any particular response based on the reproducible aspects of the neural response under normal speech listening conditions. The last three DSS components (with the lowest reproducibility from the same dataset), were similarly used to train a separate linear decoder, used as a reference to estimate baseline. In each case, the decoding procedure produces a timeseries whose similarity with the original envelope was assessed via Pearson’s *r* correlation coefficient. Each similarity score was respectively designated as reproducible (*r*_e_), and reference (*r*_f_). This referencing procedure is necessary to obtain a baseline in decoding performance since time series’ lengths varied across conditions (as a result of the different repetition rates and verses involved); otherwise there would be positive biases in *r* for shorter sequences, irrespective of underlying relationship to the stimulus.

To compute reconstruction effect sizes, each of the Pearson’s *r* pairs (reproducible versus reference activity) were transformed to Cohen’s Effect Size *q* (Cohen, 1988) by the transform 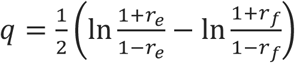. Relative effect sizes (speech vs. noise reconstruction) were computed by the fraction *q*_2_/*q*_1_ of reconstruction effect sizes given the stimulus presentation conditions above (expressed as percentages), where *q*_1_ denotes the effect size obtained from reconstructions of clean speech from neural activity following clean speech, and *q*_2_ the effect size from reconstructions of clean speech from neural activity arising from the noise probe (devoid of speech).

#### 2.5.2 Temporal response function of stimulus representation

The input-output relation between a representation *S*(*t*) of auditory stimulus input and the evoked cortical response 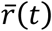 is modeled by a temporal response function (TRF). This linear model is formulated as:

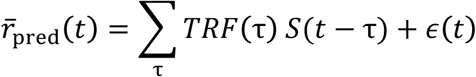

where *ϵ*(*t*) is the residual contribution to the evoked response not explained by the linear model. As stimulus representation, the envelope was extracted by taking the instantaneous amplitude of each channel’s analytic representation via the Hilbert transform (Bendat and Piersol, 2010), with sampling rates reduced to 1 KHz, transformed to dB-scale. The response was chosen to be either the first or second DSS component (fixed for each subject), according to which one produced a TRF with a more prominent M100_TRF_, a strong negative peak with ∼100 ms latency (Ding and Simon, 2012b).

#### 2.5.3 Statistical analyses

For reconstructions, one-way repeated measures ANOVA were run across the four levels: ‘Control’, and ‘Low’, ‘Medium’, and ‘High’ repetitions, in order to examine differences between their related means overall. Cortical latency of the temporal response function was determined by the M100_TRF_ latency. Peak delays with respect to control conditions were determined by cross-correlations of the TRF in the ‘Control’ versus all other repetition conditions. The resulting peak delays were then submitted to a non-parametric one-tailed two-sample Kolmogorov-Smirnov test for differences in the underlying delay populations.

## 3 Results

### 3.1 Reconstruction of missing speech from noise with context

Fixed-duration spectrally-matched static noise bursts were used to mask connected syllable/word sets within a narrated poem. Each noise probe was designed to have the same spectral composition over time as the replaced speech segment (Fig. 1A), without any supporting temporal modulations in the low-frequency (2-8 Hz) envelope (Ding and Simon, 2012a; Giraud et al., 2000). For natural speech without masking, these low-frequency fluctuations generate time-locked auditory cortical activity recorded by MEG and, given a suitable decoding model, can be used to reconstruct the envelope of the original speech signal. Such linear decoders were created to establish an optimal mapping from cortical activity to the original unmasked speech envelope. To test whether acoustic presence is a necessary condition for reconstruction of continuous speech, the listeners were exposed to extensive repetitions of some verses (each verse being a quatrain of four lines), and less frequent repetitions (or none at all) to the rest (Fig. 1B). Sentences that were maximally repeated (High repetition rate) over the hour-long session resulted in greatest relative performance in reconstruction of the envelope of the missing speech: approximately 25% of the performance for actual speech presented without any masking. Less exposure resulted in further reductions in relative performance (Medium: 21%, Low: 9%, and Control: 8%, respectively), down to the floor level in the case of masked speech with which the listener had little or no prior experience (Fig. 1C; percentages inset within each bar). Because this measure is relative to clean speech reconstruction, a measure of reconstruction from noise alone was also employed, using Cohen’s *q* to quantify the effect size. Effect sizes in reconstruction of the missing speech envelope were confirmed to display a similar pattern as with relative performance (High: 0.079 ± 0.013; Medium: 0.060 ± 0.011; Low: 0.020 ± 0.013; Control: 0.018 ± 0.008)(Fig. 1C). A one-way repeated measures ANOVA with four repetition levels was applied to determine whether decoding success of the linear model of the envelope significantly changed across conditions. Results for independent reconstructions using exclusively noise-derived *q* scores had shown that the sphericity condition was not violated (Mauchly test, *χ*^2^(5)=6.322; *p*=0.276). The subsequent ANOVA resulted in a significant main effect of repetition (*F*(3,102)=8.070; *p*<0.001). Post hoc pairwise comparisons using Bonferroni correction revealed that this increased exposure to speech significantly improved the stimulus reconstruction effect size from Control and Low repetition rate conditions to High (*p*=0.002 and *p*=0.001 respectively), and also from Control to Medium (*p*=0.008).

**Figure 1.**
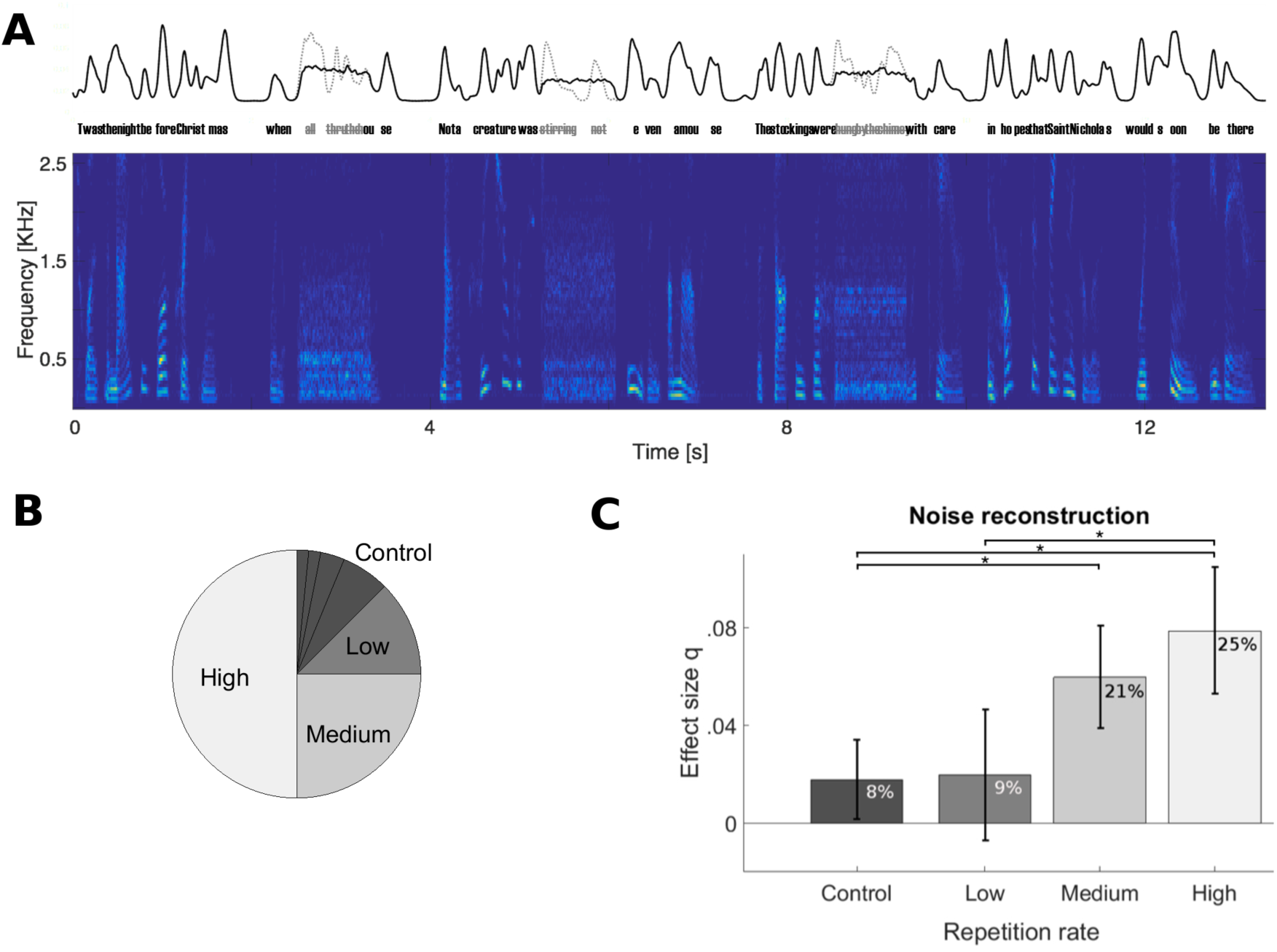
Cortical reconstruction of acoustically missing multi-word speech envelope from noise, as a function of repeated replays. (**A**) Speech material from a poem was repeatedly presented to 35 listeners, but every 4-5 s some speech was replaced with spectrally-matched noise (0.8 s duration; three instances shown in spectrogram, bottom). This manipulation removes critical temporal modulation due to the missed words, as shown by the slow envelope (top). (**B**) Some verses were presented multiple times, taking up 50%, 25%, 12.5%, etc. of all verse presentations during the hour-long MEG recording session. (**C**) The missing dynamic speech envelope could nevertheless be reconstructed from responses to the static noise that replaced the missing speech, with performance up to 25% of that obtained under clean conditions (percentages inset within each bar). This effect was not an artifact of changing contributions from clean speech reconstructions, as indicated by an alternate measure of performance, i.e., normalized with respect to independent noise-trained decoders (scale on vertical axis, right). Error bars indicate confidence intervals for the means (Bonferroni-corrected α-level).

### 3.2 Expedited auditory cortical processing of frequent natural speech replays

The temporal response function (TRF) is a functionally informative statistic, derived from a linear model, that predicts the neural response to sound stimuli, via a representation of the stimulus such as the acoustic envelope. Its characteristic peaks, and especially their polarity and latencies, are indicative of distinct neural processing stages, akin to the distinct generators of evoked responses to simple sounds such as pure tones, but directly derived from the neural processing of continuous speech (Cervantes Constantino et al., 2017; Ding and Simon, 2012a, 2012b). We examined the effect of prior exposure on the TRF’s temporal structure in general, and also for a specific peak, the M100_TRF_, occurring 100-200 ms post envelope change (Fig. 2A). When a given speech sequence was listened to repeatedly, a significant within-participant latency shift of 5.3 ± 2.2 ms earlier was observed for 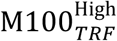 versus 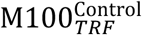 peaks (*t*(33)=2.387; *p*=0.023), indicating expedited cortical processing cortical for more familiar stimuli (Fig. 2B). Across participants, the differences between repeated (High, Medium and Low) and baseline (Control) levels, in terms of maxima in their cross-correlation functions, were shown to arise from significantly different distributions (*D*=0.294; *p*=0.043), suggesting that prior experience by repeated presentations effectively speeds up cortical processing even as early as 100 ms latency.

**Figure 2.**
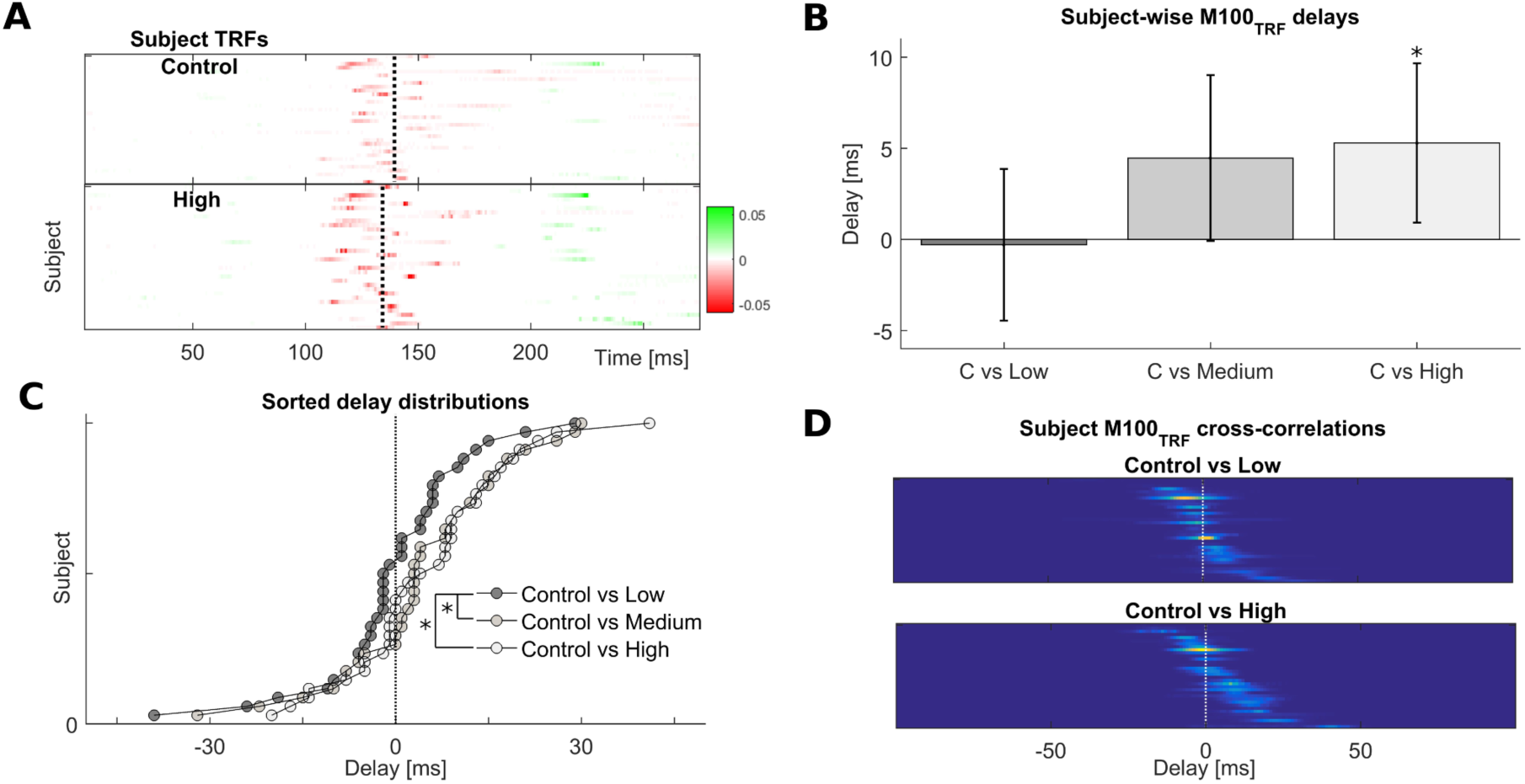
Frequent repetitions of natural speech speed-up their cortical processing. (**A**) Temporal response functions across participants reveal a common cortical processing step, referred to as the M100_TRF_, typically occurring about 100 ms after a speech envelope fluctuation (red colored features near the vertical dotted lines). (**B**) Depending on familiarity with the speech tokens, the same processing step may shift in time: processing of frequently-repeated speech occurs about 5 ms earlier than for novel or sparsely presented sentences, within subjects. (**C**) Across subjects, the distribution of relative delays is consistently biased towards positive (earlier) values for the most extreme repetition conditions. (**D**) Illustration of how shifts within subjects were obtained, by cross-correlating individual M100_TRF_ peak profiles obtained per condition in each subject.

## 4 Discussion

The phenomenon of sensory restoration relies on inference regarding elements missing from a sensory signal. The results here demonstrate that auditory cortical activity measured by MEG contains information to reconstruct the missing sequences of speech replaced by noise, provided that a listener was previously and repeatedly exposed to the missing speech. Results therefore suggest that prior experience enables access and maintenance of a detailed representation of the stimulus, in a template format compatible with the dynamical acoustic envelope; a process that may in addition be related to speed up of cortical processing time. Together, these results point to the generation of a time-locked, internally generated neural activity pattern consistent with the expected but absent sensory input. These findings complement those from related experiments investigating restoration at the phoneme-duration scale (e.g., disruptions lasting < 200 ms), which show that the acoustic presence of a specific sound pattern is not necessary for spectrogram reconstructability when speech is replaced by noise (Leonard et al., 2016) – as long as the immediate acoustic context is consistent with the restored phoneme. These results imply that the corresponding neural activity must rely on endogenous processes, possibly as top-down context-based modulations of auditory cortex populations (Petkov et al., 2007; Petkov and Sutter, 2011). The results here are consistent with the notion that this activity can be influenced by prior learning and storage of speech information, even at the level of its explicit temporal structure. Under this interpretation, enhanced listeners’ expectations about forthcoming speech tokens may predispose them to restorative encoding, in contrast to the case when contextual information is poor or insufficient, where endogenous neural dynamics may fail to adhere to or predict the missing stimulus representation. Spontaneous neural background activity known to influence perceptual processing in general, includes the ability to entrain to a complex, natural signals such as speech (Ding et al., 2013), to optimize behavioral performance of detection tasks (Henry and Obleser, 2012), or even to increase the robustness of certain auditory illusory experiences (Riecke et al., 2009).

### 4.1 Plausibility of auditory memory involvement in context effects

The auditory restoration effect investigated here may be considered part of the multimodal class of *attractive temporal context effects* (Snyder et al., 2015), a group of facilitatory mechanisms including perceptual hysteresis (Kleinschmidt et al., 2002; Schwiedrzik et al., 2014) and perceptual stabilization (Pearson and Brascamp, 2008) in the vision literature. These are considered critical for improving perceptual invariance in the face of external demands imposed by discontinuously fluctuating, broadly cluttered environments. Conceptually, this class stands opposite to that of *contrastive* temporal context effects, which are mainly suppressive, habituation or fatigue-based biases that discount neural activity after repetitions, and effectively favor perceptual alternatives for which neural activity has not yet been adapted (Schwiedrzik et al., 2014; Snyder et al., 2015). These may include semantic satiation effects, i.e., the subjective experience of increasingly meaningless words after fast and prolonged repeats (Kounios et al., 2000; Pilotti et al., 1997). Some conceptual frameworks for the organization of auditory cortical areas integrate neural coding functions with cognitive and adaptive functions such as relevance analyses of sound features, and their storage, directly in primary cortical areas (Weinberger, 2004). Storage of present connected speech sequences into sensory memory would then require retention of memory traces over the span of a few seconds, as well as past completion of stimuli resolution rendered by composite collections of features that are more efficient for long term storage (Cowan, 1984). Sensory memory has been argued to assist in the ability to restore missing fragments of a sound source, e.g. as an internal replay of the fragment during phonemic restoration (Shinn-Cunningham, 2008), and the involvement of memory-based reactivation in perceptual processes, including attention, is an area of active research (Backer and Alain, 2012, 2014; Zimmermann et al., 2016).

### 4.2 Access and format of stored auditory representations

Over the course of acoustic stimulus repetitions, attractive contextual effects may rely on implicit auditory memory, which is considered to regularly intervene in sensory and perceptual encoding (Snyder and Gregg, 2011). One such example is the improved detection of arbitrary noise structures after sequential presentations, and the time-locked potential sensory covariates of this improvement (Agus et al., 2010; Andrillon et al., 2015). Foreknowledge of acoustic features may allow listeners to adapt to a likely communication source, as demonstrated by perceptual facilitation when advance notice about the identity of a forthcoming instrument play is given (Crowder, 1989), and by preferential activation in auditory association areas specific to speaker familiarity (Birkett et al., 2007). The notion that strong expectations of a dynamic sound pattern influence the level of detail accessible in sensory representations is supported by findings of differential activation in implicit memory tasks with varying rates of sensory update: initially, short storage intervals may be associated with activation of posterior superior temporal areas, and over time, activity can be mediated by structures in inferior frontal cortex (Buchsbaum et al., 2011). Evidence from these studies is consistent with the hypothesis of transformation of memory trace representation formats, where readout from sensory buffers is at high temporal resolution under low-level representation formats, while coarser temporal resolutions may occur instead at stores that encode categorical higher-order input features (cf. Durlach and Braida, 1969; Winkler and Cowan, 2005).

### 4.3 The role of auditory imagery and related retrieval processes in listening in noise

Perceptual restoration phenomena, including phonemic restoration, may be related to auditory imagery defined as the persistence of an auditory experience without prompting by direct sensory input (Intons-Peterson, 2014). During stimulus masking, sensory imagery is postulated to involve ‘schemata’ or prior abstractions actively formed with perceptual input that become better resolved with increased familiarity, and which may remain online while an expected stimulus fails to occur (Hubbard, 2010). The implication, for methodological purposes, is that the occurrence of auditory imagery processes can be judged either by subjective reports or by using tasks hypothesized to involve imagery with reasonable probability (Hubbard, 2010). This latter approach employs familiarity of prior experience as a condition for stimuli to automatically evoke auditory imagery of original natural sound pieces (Bailes, 2007; Meyer et al., 2007). Neurally, the planum temporale is a major computational hub for which activation levels may correlate with self-reported levels of engagement with imagery, or with perceived vividness by listeners (Zatorre et al., 2009), and auditory imagery and (related) rehearsal of natural complex sounds may be subserved by auditory association cortex areas therein (Hubbard, 2010; Martin et al., 2014). There is also evidence for a dual format of representations sustained during active rehearsing, under both auditory-specific (sometimes termed ‘echoic memory’) and modality-general codes; these two coding schemes have been indicated over distinct locations each on superior temporal cortical areas, with distinct timescales as transient (< 5 s) versus sustained phases respectively (Buchsbaum et al., 2005; Meyer et al., 2007). The present data are thus consistent with a common theme in auditory retrieval processes, for which task-relevant stimuli and/or features may rely on maintenance of (re)activated domains within the sensory representational space (Kaiser, 2015). This is also supported by findings of retrieval processes in vision and hearing that involve reactivation of sensory regions active during perception (Wheeler et al., 2000), something also found with auditory verbal imagery (McGuire et al., 1996; Shergill et al., 2001), overall pointing to the notion that both involve overlapping processes (Hubbard, 2010).

### 4.4 Adaptive dynamics of speech encoding and representation during masking

The brain’s utilization of a neural model of speech input, used dynamically to infer the content of bottom-up sensory information (Pouget et al., 2013), indicates two separate but related strategies. First, the finding that cortical processing is sped up under the same circumstances that promote neural restoration of speech-coherent neural activity suggests that active, task-related endogenous processes directly optimize low-level speech processing with relevant experience. One plausible mechanism is increased excitability in a population which normally only becomes active at later stages of speech processing. Determining conditions under which this occurs may in the future provide real-time noninvasive indices of the subjective states by which a person maintains in register a template auditory pattern. Second, our results are consistent with the suggestion that auditory ‘image’ formation entails activity consistent with that elicited by original sound input (Janata, 2001; Martin et al., 2017), where preservation of the temporal acuity (and related properties) of the original stimulus may deteriorate depending on factors such as context and experience (Janata and Paroo, 2006). The latter appears related to the different success rates in reconstruction of missing speech found here, which decreased for increasingly unfamiliar stimuli. A need for frequent “refreshing” then echoes the auditory memory reactivation hypothesis where storage of individual sound features is embedded in the context of those neighboring patterns and sequences representable by the auditory system as regularities. Reactivation here denotes the automatic process where variable sound input is matched to constancies extracted previously; likelihood of storage is then increased by proximity between a prior rule and current update tokens (Winkler and Cowan, 2005). This description, originating from oddball sequence studies, can be considered to apply in the present study across its verse stimulus structure: e.g., dynamic acoustic features of speech preceding a masker may serve as referents for a listener, enabling the process of translation of verse regularities learned and represented over the course of the experiment, into specific values in the same feature format (Winkler and Cowan, 2005). While this does not preclude additional dynamic stimulus features also contributing, including higher-order linguistic elements (e.g. Di Liberto et al., 2015; Kayser et al., 2015; Näätänen and Winkler, 1999; Wassenhove and Schroeder, 2012), the suggestion that a key neural property of natural sound encoding is via temporally-based acoustic representations is underscored by their active maintenance during noise gaps, based on prior experience.

## Conflict of Interest

The authors declare that the research was conducted in the absence of any commercial or financial relationships that could be construed as a potential conflict of interest.

## Funding

This study was funded by the National Institutes of Health (R01-DC-014085).

## Acknowledgments

We thank Anna Namyst for excellent technical assistance.

## Data availability

All relevant data are available to all interested parties in a public repository at <http://hdl.handle.net/1903/20259>.

